# Biophysical analysis reveals autophosphorylation as an important negative regulator of LRRK2 dimerization

**DOI:** 10.1101/2023.08.11.549911

**Authors:** Giambattista Guaitoli, Xiaojuan Zhang, Francesca Saitta, Pasquale Miglionico, Laura M. Silbermann, Franz Y. Ho, Felix von Zweydorf, Marco Signorelli, Kasia Tych, Dimitrios Fessas, Francesco Raimondi, Arjan Kortholt, Christian Johannes Gloeckner

**Affiliations:** German Centre for Neurodegenerative Diseases, 72076 Tübingen, Germany; Department of Cell Biochemistry, University of Groningen, Groningen 9747 AG, The Netherlands; Department of Food, Environmental and Nutritional Sciences, University of Milan, 20133 Milan, Italy; Scuola Normale Superiore, 56126 Pisa, Italy; Groningen Biomolecular Sciences and Biotechnology Institute, University of Groningen, Groningen 9747 AG, The Netherlands; YETEM-Innovative Technologies Application and Research Centre, Suleyman Demirel University, Isparta 32260, Turkey; Core Facility for Medical Bioanalytics, Institute for Ophthalmic Research, Center for Ophthalmology, University of Tübingen, Tübingen, Germany

## Abstract

Leucine-rich repeat kinase 2 (LRRK2) is a large, multi-domain protein which is associated with Parkinson’s disease. Although high-resolution structures of LRRK2 are available, little is known about the complex dynamics behind the inter-domain regulation of LRRK2 and its perturbation by pathogenic variants. Previous studies have demonstrated that LRRK2 goes through an oligomerization cycle at the membrane, however it remains unclear in which form it exerts its kinase activity. Moreover, the LRRK2 monomer-dimer equilibrium and associated functional implications at a molecular level also need further investigation. In the present work, we used a multi-faceted approach to better understand LRRK2 oligomerization and suggest a functional model of how LRRK2 interacts with its substrates. To this end, we combined nano differential scanning calorimetry and mass photometry with molecular modelling. The thermal analysis resulted in a multistep denaturation profile, elucidating novel insights into the composite structural organization of the multi-domain protein LRRK2. Furthermore, LRRK2 shows a remarkable thermal stability, confirming its oligomeric nature. By using mass photometry, we could observe a monomer-dimer equilibrium which is altered by R1441G, a pathogenic variant within the Roc-COR interface. Most importantly, we could demonstrate that autophosphorylation induces LRRK2 monomerization, indicating a novel intramolecular feedback mechanism. Finally, we investigated the interaction of LRRK2 with its substrate, RAB10 by integrative computational modelling. The resulting models suggest that the monomeric form of LRRK2 is the favored protein conformation for the interaction with its substrate, leading to an increasing interest in the monomer-dimer equilibrium as a possible intervention point for the pathology.

## Introduction

Parkinson’s disease (PD) is the second most common age-related neurodegenerative disease and is clinically characterized by movement impairments, bradykinesia, rigidity and resting tremors (Fahn, 2003). Although most forms of the disease are sporadic (idiopathic PD), familial forms of PD exist (Gasser, 2009). Missense variants in the Leucine-Rich Repeat Kinase 2 (*LRRK2*) gene have been identified to cause familial forms of PD but risk variants within its genomic locus (PARK8) also play a role in the etiology of iPD (Simon-Sanchez et al., 2009; Zimprich et al., 2004). Furthermore, a recent study suggests that, independent of mutations, increased LRRK2 activity plays a role in iPD (Di Maio et al., 2018). LRRK2 encodes a large ubiquitously expressed multi-domain protein of 286-kDa and 2527 amino acids that exerts both GTPase and kinase activities (Biosa et al., 2013; Gloeckner et al., 2006; West et al., 2005). LRRK2 is involved in multiple cellular processes including protein scaffolding, regulation of cytoskeletal dynamics and vesicle sorting but has also been identified as a regulator in innate immunity (Ahmadi Rastegar and Dzamko, 2020; Gloeckner and Porras, 2020; Pellegrini et al., 2017; Roosen and Cookson, 2016).

Various PD-associated mutations in LRRK2 have been shown to augment kinase activity leading to an increased auto-phosphorylation as well as phosphorylation of Rab proteins, the recently identified physiological LRRK2 substrates (Gloeckner et al., 2006; Greggio et al., 2006; Jaleel et al., 2007; Steger et al., 2016; West et al., 2005). Furthermore, pathogenic LRRK2 mutations within the Roc domain have been associated with decreased LRRK2 GTPase activity.

The protein belongs to the Roco-protein family of G-proteins. Members of this family have in common a Ras of complex proteins (Roc) G-domain and an adjacent conserved C-terminal of Roc (COR) acting as dimerization domain (Gilsbach and Kortholt, 2014). Besides its enzymatic core, LRRK2 contains four predicted solenoid domains, commonly involved in protein–protein interactions. These domains include the N-terminal Ankyrin, Armadillo, and namesake Leucine-Rich Repeat (LRR) domains, along with a C-terminal WD40 domain (Mills et al., 2012).

Membrane localization plays an important role in the oligomerization and subsequent activation of LRRK2 (Berger et al., 2010) where it phosphorylates a specific subset of proteins (Gomez et al., 2019). In addition, LRRK2 seems to be predominantly monomeric in the cytosol while forming oligomers at the membrane (James et al., 2012). Structural models as well as high-resolution structures, which became available recently, suggest that the LRRK2 N-terminus as well as the C-terminal WD40 domain may also be involved in auto-regulatory processes, likely keeping LRRK2 in an auto-inhibited state and, besides the Roc-COR module, may also contribute to the LRRK2 oligomerization (Deniston et al., 2020; Guaitoli et al., 2016; Watanabe et al., 2020).

Available high-resolution structures, together with biochemical studies already gave valuable insight into the local interaction of different domains and helped to identify first regulatory mechanisms of the LRRK2 kinase activity (Deniston et al., 2020; Taylor et al., 2020; Zhu et al., 2022). Activity-dependent local transitions within the LRR-Roc linker region have also been recently identified by molecular dynamics (MD) simulations (Weng et al., 2023). However, the dynamic process of LRRK2 regulation by conformational and oligomerization changes remains largely unknown.

In the present work, differential scanning calorimetry (nDSC) of the purified human full-length LRRK2 dimer protein gave valuable insights of the conformational stability of the apo-protein, as well as of the protein phospho-states thermal stability. Furthermore, mass photometry (MP) measurements allowed the determination of the kinetics of the LRRK2 monomer-dimer and revealed functional roles of auto-phosphorylation in the oligomerization process.

Finally, Crosslinking mass spectrometry (CX-MS)-driven integrative models of the LRRK2:Rab10 complex suggest that the monomer is the favorite state allowing kinase-substrate interactions. Unbiased AF-multimer predictions converge on predicting similar LRRK2-KIN-RAB10 interfaces, which are consistent with several of the XL as well as other biochemical evidences (e.g. RAB10 T73 phospho-site).

## Results

### LRRK2 thermal stability

To determine the stability of the LRRK2 dimer conformation, the full-length LRRK2 thermal denaturation was assessed by nano-DSC (Fig. 1, blue profile). For this purpose, recombinant Strep-FLAG (SF)-tagged full-length LRRK2 was purified from HEK293T cells (Supplementary Fig. 1). The observed profile shows a very broad shouldered and asymmetric endothermic signal corresponding at the thermal protein denaturation process. However, aggregation phenomena were observed at high temperatures (see Material and methods section), preventing us from performing a straight equilibrium thermodynamic analysis in absolute enthalpic and entropic terms. Nonetheless, the intensive properties, i.e., both the denaturation temperature range and the calorimetric profile, are still highly informative and permitted at least a qualitative assessment of the overall denaturation mechanism (see Supplementary Material for details).

**Figure 1.**
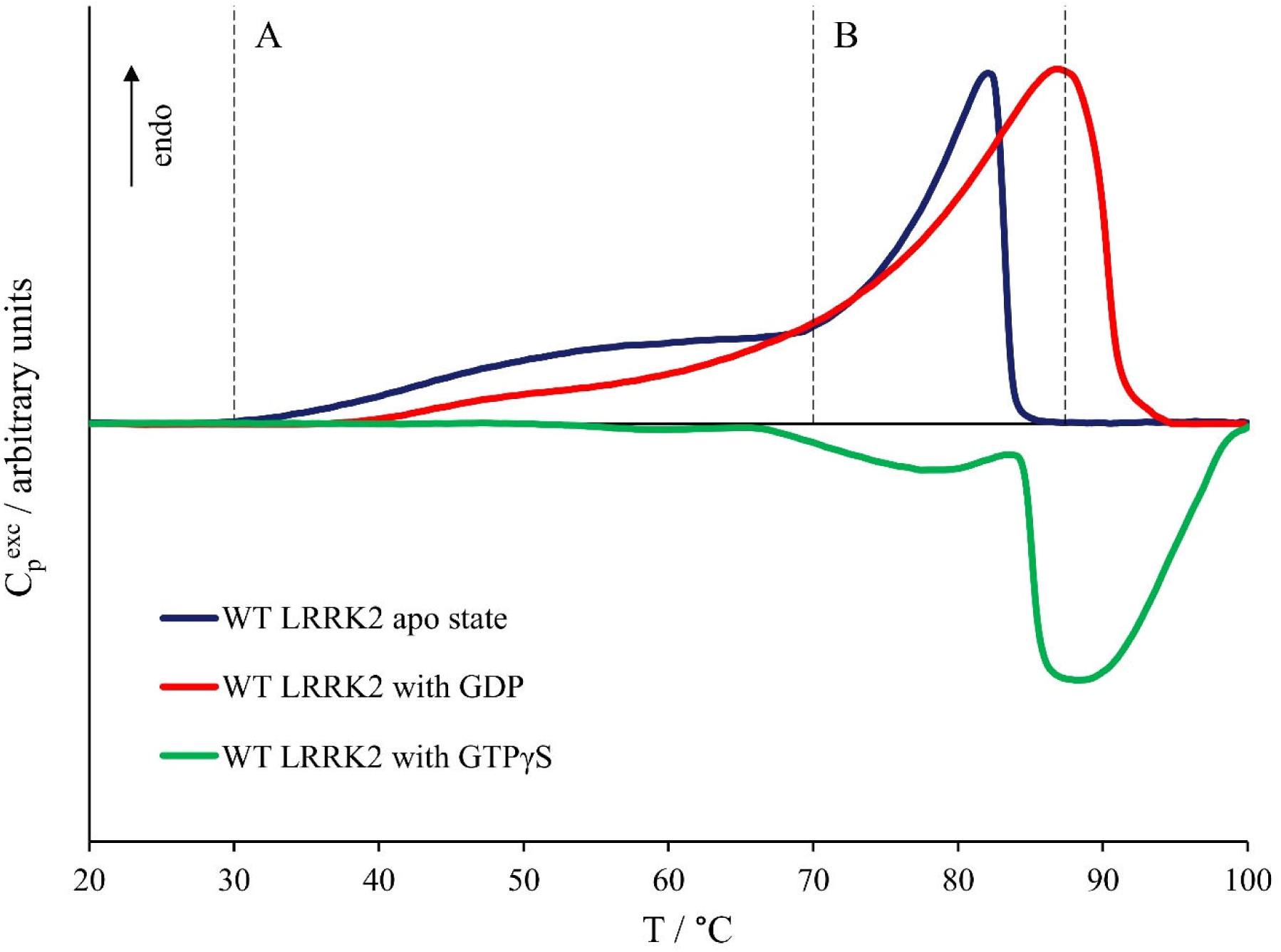
Nano-DSC thermograms obtained for the purified full-length LRRK2 apo state (blue trace) and in the presence of either GDP (red trace) or GTPγS (green trace).

Indeed, the first part of the thermogram (region A, Fig. 1) indicates the presence of several thermodynamic domains that exhibit low stabilities, which are compatible with relatively less compact protein structural domains (Fessas et al., 2001). In general, there is not straight correlation between thermodynamic and structural domains but, in many cases, thermal stability profile has been successfully used as a structure-correlated fingerprint to assess protein structural peculiarities (Durowoju et al., 2017; Fessas et al., 2001; Fessas et al., 2007). Being aware of the uncertainties and the need of further confirmation, we highlight that these preliminary indications are in line with the data obtained from the recently published Thermal Shift Assay data for a LRRK2 Kinase-WD40 domain construct (Schmidt et al., 2021). According to this analysis, the denaturation temperatures for Kinase and WD40 domains were measured to be 45°C and 56°C, respectively, collocating them in the low-temperature portion of the overall protein thermal denaturation profile. By contrast, the thermogram portion over 70°C (region B, Fig. 1) reveals a calorimetric profile that is typical of a denaturation mechanism that complains a dissociation process (Ausili et al., 2013; Fessas et al., 2001) concomitant with the denaturation of a rather stable structures as indicated by the peak size if compared to the previous domains’ thermal denaturation signal. A more detailed description of the thermodynamic analysis of the LRRK2 apo-state can be found in the supplementary information.

In order to assess the effects of defined G-nucleotide states on the full-length LRRK2 thermal stability, nDSC measurements were performed by alternatively loading the protein with GDP or GTPγS, a non-hydrolysable GTP analogue. Fig. 1 shows the thermal denaturation profile of the GDP-loaded full-length LRRK2 (red profile). We observed again a very broad endothermic and asymmetric thermogram corresponding to the thermal denaturation of the protein, in first approximation similar to the profile observed for LRRK2 in its apo-state, indicating a similar denaturation mechanism. Unfortunately, post-denaturation aggregation phenomena were observed also in this case, though to a minor extent. Accordingly, we limit at comparing the position and shape of the calorimetric profiles being the differences between the two conditions clearly visible. Indeed, the low-temperature portion of the thermogram results to be reduced with respect to the second part, which, by contrast, seems to be enhanced and shifted towards higher temperatures (*T_max_*at about 86°C).

Conversely, as far as the effect of GTPγS on LRRK2 is concerned, the nDSC thermogram shown in Fig. 1 (green profile) still discloses some endothermic thermodynamic steps in the range between 40 and 80°C, which unfortunately cannot be clearly discriminated due to the predominant aggregation and precipitation exothermic phenomena preventing any further analysis at these experimental conditions.

Together our data show that LRRK2 forms a remarkable thermal stable dimer. Furthermore, the presence of G-nucleotide affects the protein overall stability, GDP-bound LRRK2 shows increased stability, while in contrast GTP-bound LRRK2 shows strongly reduced thermal stability considering the early post-denaturation aggregation phenomena.

### G-nucleotide dependency of LRRK2 conformational states

Structural and biochemical studies have shown that LRRK2 exist both as a monomer and dimer (Deniston et al., 2020; Taylor et al., 2020; Zhu et al., 2022), however none of the methods used so far allowed determining the kinetics and mechanism of that regulates LRRK2 monomer-dimerization. To characterize the monomer-dimer equilibrium, we subjected LRRK2 to Blue-Native PAGE analysis and MP. The oligomeric state of the bacterial LRRK2 orthologue CtRoco correlated to defined G-nucleotide states, with a monomeric GTP and dimeric GDP-bound conformation (Deyaert et al., 2017). However, Blue-Native Page analysis with purified WT LRRK2 revealed that, regardless of the nucleotide binding state, a single band corresponding to a dimeric protein (Fig. 2, Supplementary Figs. 1C, D). However, Blue-Native PAGE requires a considerable high sample concentration (approx. 2µM) to allow the detection of LRRK2 by colloidal Coomassie, which might force its dimerization. In contrast, MP, an interferometric scattering microscopy, allows to determine the molecular mass distribution within a protein sample at low concentrations (100pM to 100nM) and minimal volumes (Young et al., 2018). Confirming our previous observations, the analysis showed two distinct particle contrasts, which transformed to molecular weights corresponding to the expected weights of the LRRK2 monomer and dimer, respectively. At the measured concentration, the protein was mainly in the monomer state (Supplementary Materials). Subsequently, we measured LRRK2 conformations under different nucleotide states (non-hydrolysable GTP analogues and GDP state) as well as concentrations (25, 50, 75, and 100nM). Interestingly, the concentration-dependent changes in the monomer/dimer ratio from MP allowed an estimation of the dissociation constant (K_d_) about 200nM for LRRK2 dimers. However, no significant differences were observed between GDP and GTP-analogue bound states of LRRK2 at the concentrations tested with this technique (Fig. 3A).

**Figure 2.**
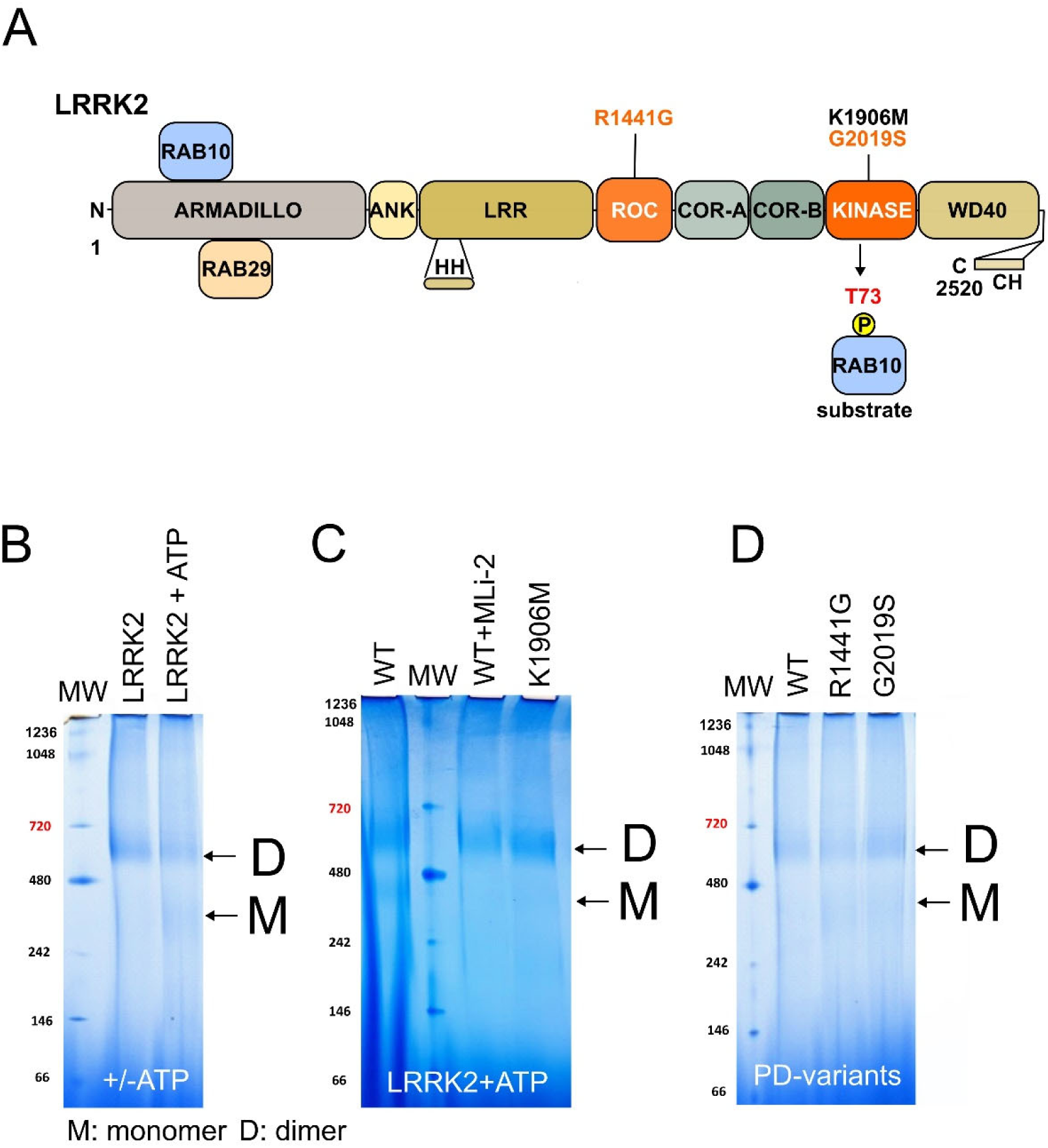
(A) Overview of the LRRK2 domains structure. Armadillo: Armadillo domain, ANK: Ankyrin domain, LRR: Leucine-rich repeats, ROC: <u>R</u>as <u>o</u>f <u>c</u>omplex proteins G domain, COR (A/B): <u>C</u>-terminal <u>o</u>f <u>R</u>oc, Kinase: kinase domain, WD40: 7-bladed WD40 propeller domain., HH: hinge helix, CH: C-terminal helix. PD variants used in this study are indicated in orange, the kinase-dead is indicated in black. N-terminal binding sites of RAB10 and RAB29 are indicated. (B-D) Blue native analysis. (B) ATP pre-incubation induced LRRK2 monomerization. (C) ATP induced LRRK2 monomerization is blocked by the LRRK2 specific MLi-2 inhibitor or a kinase-dead variant. (D) Effect of PD variants on the monomer/dimer ratio.

**Figure 3:**
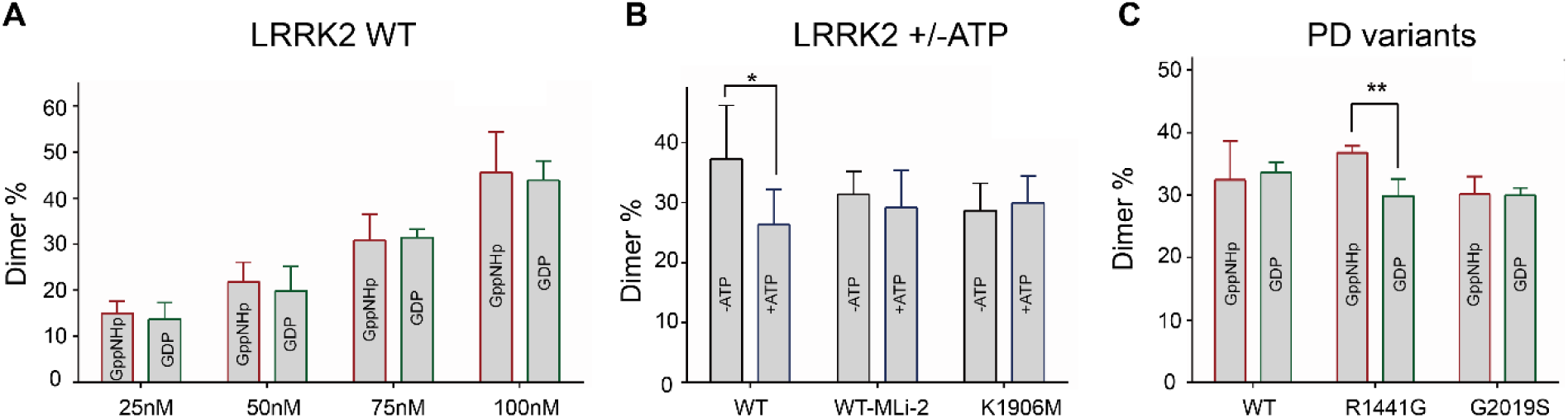
Analysis of the LRRK2 monomer/dimer (M/D) equilibrium by mass photometry. (A) M/D equilibrium at different LRRK2 concentration in dependence of the G-nucleotide (GppNHp or GDP). (B) Effect of ATP-preincubation on the M/D ratio. (C) Impact of PD variant on the M/D equilibrium in dependence of the G-nucleotide (GppNHp or GDP). Statistical analysis (N=3) was performed by a T-test. Significant changes are indicated by Asterisks: *P<0.05; **P<0.01.

Together, our data show there is monomer and dimer state of LRRK2, *in vitro*. While the nucleotide might make some contributions to LRRK2 stability, we did not observe a nucleotide-dependent conformation as seen in the homogenous bacterial protein (Deyaert et al., 2017).

### Autophosphorylation induces LRRK2 monomerization

Recently we have revealed a feed-back regulatory mechanism of the kinase domain on GTPase activity. In that study, we could show that the kinetical properties of LRRK2-mediated GTP hydrolysis are negatively regulated by auto-phosphorylation (Gilsbach et al., 2023). To assess if this negative feedback is mediated via LRRK2 monomerization-dimerization, we auto-phosphorylated the purified protein under conditions allowing sufficient phospho-transfer (Gloeckner et al., 2010) and analyzed its oligomeric state. In the presence of ATP, Blue-Native PAGE analysis revealed in addition to the band corresponding to the LRRK2 dimer, a second band at the expected molecular weight of the monomer (approx. 300kDa) (Fig. 2B). Consistently, mass photometry demonstrated that LRRK2 pre-incubated with ATP shows significantly less dimerization (Fig. 3B). To confirm that the decrease in dimeric protein was indeed due to the ATP-induced auto-phosphorylation and not due to occupancy of the ATP binding site, we repeated the experiments in presence of the LRRK2-specific kinase inhibitor MLi2. No difference was observed between ATP-treated or untreated LRRK2 in presence of MLi-2, confirming that the observed shift in the monomer/dimer equilibrium was indeed the result of auto-phosphorylation. Similarly, no monomerization could be induced by ATP-treatment for the kinase-dead mutant LRRK2 K1906M (Fig. 3B).

In conclusion, these data implicate that the LRRK2 autophosphorylation regulates the monomer/dimer equilibrium.

### Impact of PD mutations on the monomer-dimer cycle

To investigate whether disease-associated variants affect LRRK2 function by disturbing the monomer-dimer cycle, we compared the conformational states of different PD variants located in the Roc domain (R1441G) as well as in the kinase domain (G2019S) with the LRRK2 wild-type in dependence of the nucleotide state. In a Blue-Native PAGE analysis no clear defect of particular PD variants on monomer-dimer equilibrium were observed (Supplementary Fig. 1C). The mutant proteins, at the high concentration required for the experiment, clearly show a predominant dimeric form, though a faded signal correspondent to the monomer can maybe be observed (Fig. 2D) Next, MP profiles were determined using LRRK2 protein purified in the presence of different nucleotides at a fixed concentration (75nM). Among the variants tested, only the R1441G mutant showed a nucleotide-dependent shift in the monomer/dimer equilibrium. In fact, a higher degree of dimerization was found for GppNHp-bound LRRK2 compared to the GDP-bound state. In contrast, there was no nucleotide-dependent effect observed for G2019S, suggesting that the underlying PD mutations located in different domains might feature distinct molecular pathomechanisms (Fig. 3C).

### Integrative modelling suggests preferential binding of RAB10 to the LRRK2 monomer

In addition to its relevance in GTPase activity, the monomer-dimer equilibrium may also play a role in the activation as well as the interaction pattern of LRRK2.

To better understand how LRRK2 physically interacts with its effectors, we followed an integrative computational modelling approach. For the initial assessment of conformational changes upon Rab binding, we employed a crosslinking approach combined with mass spectrometric mapping of the linked lysine residues which we have previously used for building a model of dimeric LRRK2 as well as to map binding sites of LRRK2-specific nanobodies (Guaitoli et al., 2016; Singh et al., 2022). Recombinant LRRK2 as well as the full-length Rab proteins were purified from HEK293T cells (Supplementary Fig. 1). When comparing the resulting crosslinking datasets of different experimental conditions generated by the CID-cleavable crosslinker DSSO, we could already draw two major conclusions: Crosslinks between RAB10 and LRRK2 were only obtained for GTP-loaded LRRK2 and in the presence of RAB29. The affinity of Rabs to LRRK2 has been previously shown to be in the lower µM range (Vides et al., 2022). To ensure binding the protein concentration has been selected, accordingly. The final LRRK2 concertation was adjusted to 1-2µM. The Rab proteins were mixed with LRRK2 in a 4-fold molecular access. While crosslinks were seen between RAB10 and Rab29, no crosslinks were seen between LRRK2 and Rab29. Multiple binding sites for Rab proteins have been described in the N-terminal Armadillo domain, which we previously found not to be well accessible to crosslinking, likely due to its compact folding (Guaitoli et al., 2016; Steger et al., 2017; Vides et al., 2022; Waschbusch et al., 2014). As Rabs are not known to form direct protein:protein complexes while interaction with LRRK2 at micromolar affinity (Vides et al., 2022), the crosslinks between RAB10 and RAB29 might be the result of constraints provided by LRRK2. In the modelling approach, we limited the analysis to the LRRK2:RAB10 complexes as we only obtained crosslinks between LRRK2 and RAB10 passing the significance filters of the analysis software. When comparing conditions +/- RAB10 we observed a re-arrangement of the intramolecular crosslinks of LRRK2, demonstrating that a major conformational change is necessary to accommodate the substrate in proximity to the enzymatic core. In details, RAB10 binding is concomitant with a loss of XL-mediated contacts mediated by the ankyrin (ANK) domain with both the N-term domains such as armadillo (ARM), LRR and ANK-LRR linker, as well as with the RCKW, in particular KIN and WD40 domains (Figs. 4A, B). At the same time, ANK replaces LRR in interaction with the COR domain. Further support for this idea comes from contact analysis of the available LRRK2 structures, by aggregating the identified contacts at the domain level. In the APO cryo-EM structures (PDB:7LHW), the RCKW is engaged in multiple contacts with N-terminal domains (Supplementary Figs. 2, 3A). In contrast, Alphafold2 models of full-length, monomeric LRRK2 favor the kinase domain in an active conformation and show a less compact folding (i.e.: AF-Q5S007-F1), which also misses potentially inhibitory contacts between the hinge helix located between the N-terminal repeats and the C-terminal WD40 domain, as visible in the recently published high resolution full-length structures (42, 46) (Supplementary Figs. 2, 3B, 4).

**Figure 4:**
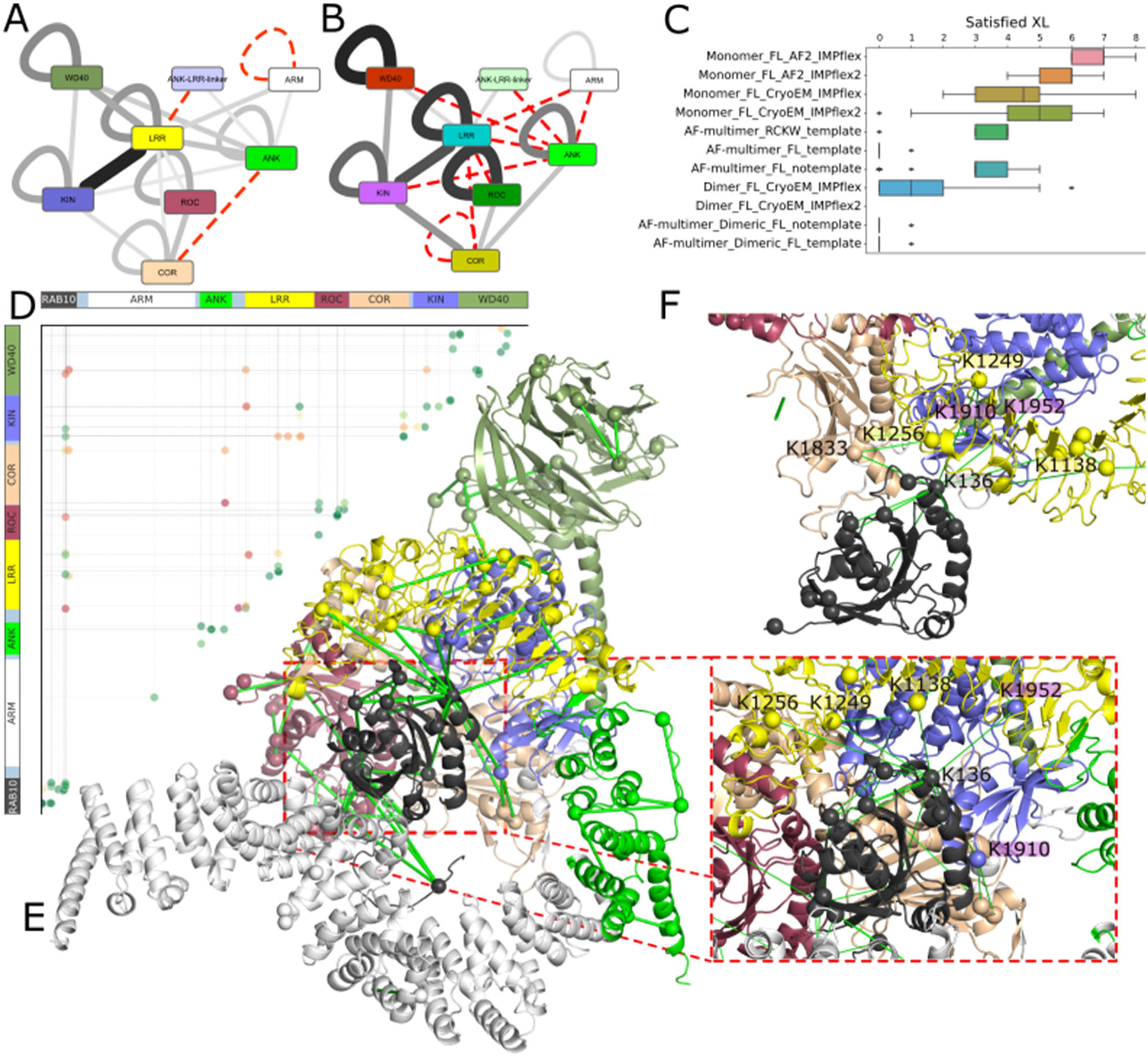
Integrative modeling of LRRK2-RAB10 complexes with CX-MS. Domain-level network representation of CX-MS data referred to LRRK2 in the (A) APO and (B) RAB10-bound states. Nodes correspond to domains, continuous edges indicate the presence of CX-MS between two domains, whose thickness is proportional to the number of XLs, dashed red edges indicate domain XL not seen in that state but observed in the other; (C) boxplot statistic of the number of satisfied cross-links for each prediction runs. For IMP simulations, only the results obtained on the complexes from the best clusters are shown. For AF-multimer predictions, the results from the top 5 ranked models are shown. Boxplots show the median as the center and first and third quartiles as bounds of the box; the whiskers extend to the last data point within 1.5 times the interquartile range (IQR) from the box’s boundaries; (D) Bubble plot showing the CX-MS connectivity in the best model predicted via AF-multimer (with no template). Greenish dots represent XLs whose connected residue lay at a distance smaller than 35Å in the AF-multimer top complex, with darker green corresponding to closer distances, while reddish dots represent XLs whose connected residue lay at a distance greater than 35Å, with darker red corresponding to larger distances; (E) cartoon representation of the best ranked model of LRRK2-RAB10 complex predicted by AF-multimer, with CX-MS related distances below 35Å represented as green continuous dashes (left); zoomed view of the LRRK2-RAB10 interface (right); (F) zoomed view of the LRRK2-RAB10 interface of the best model predicted by IMP.

Next, we used available structures to model the LRRK2:RAB10 complex through CX-MS driven integrative modeling through Integrative Modeling Platform (IMP) (see Material and Methods). We considered as input for the calculations both monomeric (experimental and AF2 predicted, see Methods), as well as available dimeric structures of LRRK2. By superimposing available APO and AF2-predicted monomeric structures, we could identify two flexible linker regions, one between the LRR and Roc domain (1328-1329) and a second between CORA and the CORB sub domains (1657-1667). We then carried out integrative modeling by considering LRRK2 and RAB10 as rigid bodies and allowing flexibility on either the LRR-ROC linker alone or in combination with the CORA-CORB linker. Assessment of satisfied XLs formed between RAB10 and LRRK2 on top-scoring, most representative models, revealed that best results are achieved when only treating the LRR-ROC linker region as flexible (Fig. 4C). This finding is in well agreement with a molecular dynamics (MD) analysis performed recently, proposing an ordered to disordered transition of the linker region (Weng et al., 2023). Integrative modeling simulations also revealed that monomeric input structures, particularly the AF2 model, achieved maximum CX-MS satisfaction, while lowest agreement between the XL data set and the models was obtained when using dimeric full-length LRRK2 structures (Myasnikov et al., 2021) as input (Fig. 4C).

As an independent approach, we also predicted RAB10 complexes by using AlphaFold-multimer, by considering either monomeric or dimeric LRRK2 states (see Methods). We validated the predicted models by mapping CX-MS and assessing their spatial satisfaction in the structural models. Also in this case, we found that the predictions with the monomeric LRRK2 led to a higher number of satisfied crosslinks than the dimeric state (Figs. 4C-E), suggesting that the former state is most likely responsible of mediating the interaction with the substrate. Intriguingly, the best complexes between monomeric LRRK2 and RAB10 predicted by AF-multimer involve the same RAB10 interface, mediating CX-MS through Lys 136 on α4 helix, in addition to contacts between RAB10 switch regions with the ARM domain (Fig. 4E). Among the alternative models predicted by AF-multimer, we also found one where the RAB10’s T73 is very close to the KIN catalytic site (Supplementary Fig. 3A). A broader modelling approach for different LRRK2:RAB complexes based on interactions curated in the InAct database is shown in Supplementary Fig. 2C. In agreement with previous studies, multiple Rab binding sites are detected in the Armadillo domain (Dhekne et al., 2023; Vides et al., 2022). However, district contact patterns with the enzymatic core (Roc-COR-Kinase) of LRRK2 have also been detected.

In particular, we found that monomeric LRRK2 is subjected to a large conformational change of both the ROC and COR domains, which are roto-translated with respect to the FL, monomeric LRRK2 APO state (Figs. 4D, 5A). The Roc-COR conformational change is propagated via the LRR-Roc linker region to the N-terminus (LRR, ANK and ARM), which is maximally displaced to accommodate the positioning of the RAB10 substrate, which on one side is docked on the ARM domain, and in the other it forms an interface with the KIN domain. More in details, we analyzed the hinge and screw axes of the conformational change required to bind RAB10 via established protocols (e.g. DynDom (Veevers and Hayward, 2019)), which highlighted a multi-step conformational change happening through multiple screw axes (Figs. 5A). One is located at the KIN-COR interface, allowing a rigid-body-like roto-translation of the Roc-COR toward the WD40-KIN (Figs. 5A-C). Another one is centered on the α0 helix linking the Roc and LRR, describing a rotation of the LRR, to allow the large displacement of the N-term (Figs. 5A, D-E). A third axis is found at the border between the ANK and ARM domain, and it is instrumental in presenting RAB10 in close proximity to the KIN domain (Supplementary Fig. 3). The AF-multimer predicted complexes between RAB10 and dimeric-LRRK2 never display such a large conformational displacement of the Roc-COR as well as the N-term domains, while RAB10 frequently docked on just the ARM domain. This is likely due to the constraint exerted by the dimeric interface, which prevents ROC-COR to rearrange and unleash the large conformational displacement of the N-term, as observed in the monomeric LRRK2 complex (Figs. 5A).

**Figure 5:**
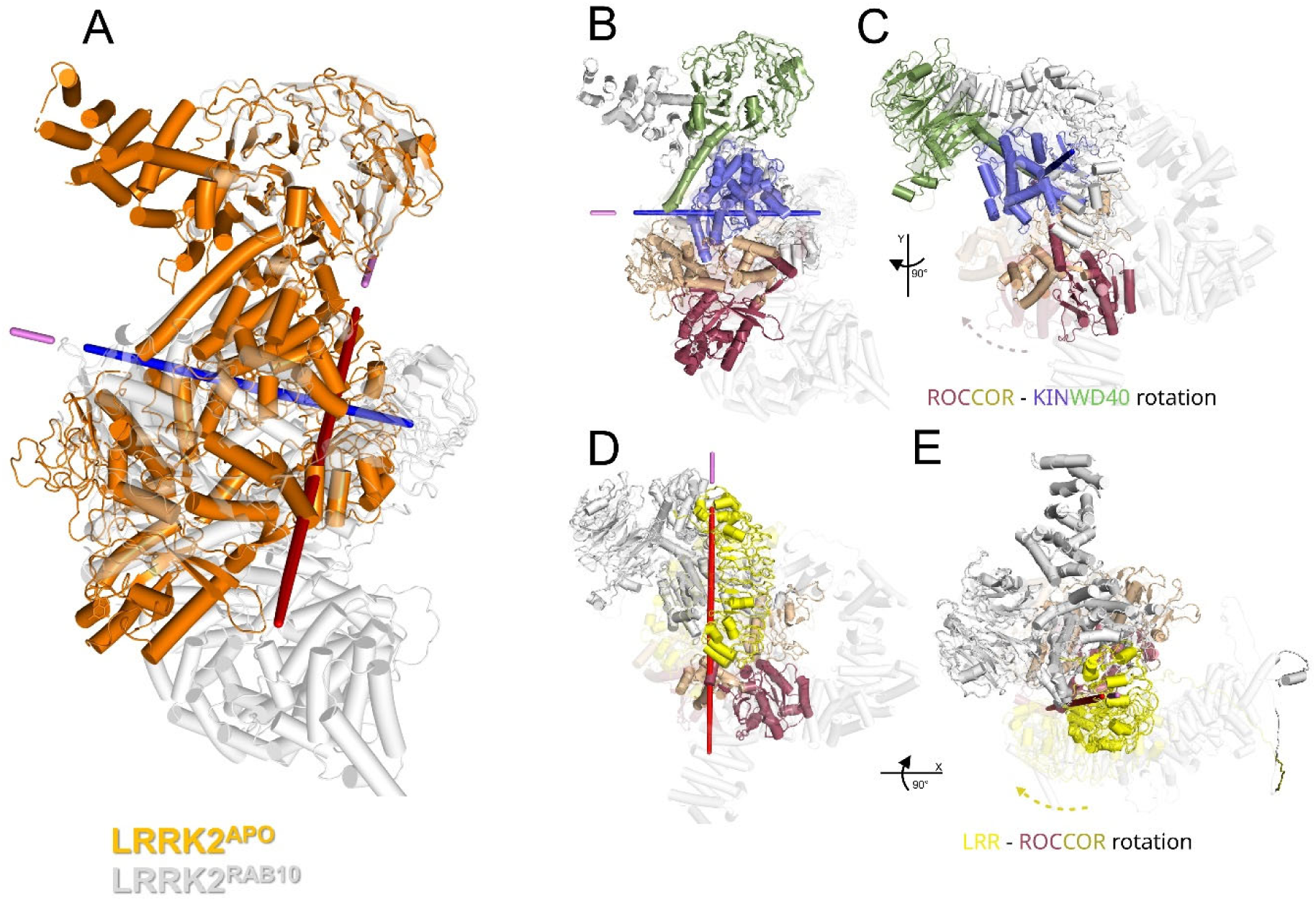
Major rearrangements are necessary to accommodate Rab10 docking on LRRK2. (A) superimposition of the LRRK2 structure in the APO state (orange; PDB 7LHW) with the structure predicted by AF-multimer in complex with RAB10 (gray). Blue and red axes indicate the main conformational changes, identified through DynDom, occurring between the ROC-COR with KIN-WD40 and LRR, respectively; (B) superimposition of the LRRK2 structure in the APO state (PDB:7LHW) with the structure predicted by AF-multimer in complex with RAB10 (gray). Blue axis indicates the conformational change identified between the ROC-COR and KIN-WD40 domains. Moving domains are colored in the APO state; (C) 90° rotated (y axis) view of the conformational change in (B); (D) Superimposition of the LRRK2 structure in the APO state (PDB:7LHW) with the structure predicted by AF-multimer in complex with RAB10 (gray). Red axis indicates the conformational change identified between the ROC-COR and LRR domains. Moving domains are colored in the APO state; (E) 90° rotated (x axis) view of the conformational change in (D).

In conclusion, the integrative computational analysis suggests that LRRK2 binds its substrate RAB10 preferably in its monomeric form.

## Discussion

Although LRRK2 is considered as one of the most promising targets for PD treatment and thus object of intense research, in particular its activation and regulatory dynamics at a molecular level as well as the impact of PD variants on these mechanisms is yet to be determined at a more detail. Roco proteins, including LRRK2, have been shown to undergo an oligomerization cycle which is considered to be a central mechanism of switch between active and inactive states (Wauters et al., 2019). For this reason, a better understanding of the oligomerization dynamics might give additional insight into the molecular pathomechanisms associated with LRRK2-PD. We used a multifaceted approach, combining biophysical and biochemical analysis with computational modelling to investigate LRRK2 oligomerization, which lead to the identification of a novel feedback mechanism driven by autophosphorylation. In addition, integrative computational models allowed new conclusions about how LRRK2 interacts with its substrates and the role of oligomerization in this process.

In the present work, we determined the thermal denaturation profiles of human LRRK2 protein. The analysis of large multidomain proteins by nDSC is generally challenging and, to our knowledge, has not done before for full-length LRRK2. By providing a detailed analysis of such profiles we could confirm the dimeric nature of purified LRRK2 and observe that its dimer has a remarkable thermal stability indicated by a melting temperature of approximately 85°C. GDP binding to purified full-length LRRK2 induces a different thermal profile between 40 to 60 °C. Interestingly, the profile in this temperature range correlates with thermal data previously determined for the LRRK2 Kinase-WD40 module (Deniston et al., 2020). Furthermore, a slight increase in dimer stability can also be observed, indicated by a slight shift of a characteristic large endothermic peak towards a higher melting temperature. However, in presence of GTPγS, a non-hydrolysable GTP analogue, the observed exothermic profile reflects an extensive structural change that leads to unstable and aggregated material. Taken together, this shows that, dependent on the nucleotide binding, the stability of the LRRK2 changes in different ways.

By mass photometry, an optical approach to determine molecular masses in solution, we could for the first time estimate a dissociation constant of around 200nM which is one order of magnitude lower compared to GDP-bound *Chlorobium tepidum* (*Ct*) Roco, a bacterial Roco protein (Deyaert et al., 2017). The observed oligomerization of *Ct*Roco is nucleotide dependent. However, based on the thermodynamic data, a G-nucleotide-dependent oligomerization process, similar to what has been seen for the bacterial homologues, has not been observed. None of the used investigation methods neither nDSC nor mass photometry could give a clear indication of a GTP-driven monomerization. However, we cannot exclude that GTP is inducing release of the LRRK2 RocCOR dimer interface, but that other domains keep LRRK2 in a dimeric conformation.

In contrast, by analyzing the most relevant PD-associated variants segregating with the disease, we could demonstrate that a variant within the Roc-COR interface, R1441G, shows a higher extent of monomerization when bound to a non-hydrolysable GTP analogue, indicating that indeed a perturbed monomer-dimer cycle could be a relevant pathomechanism.

Furthermore, and rather unexpected, we could identify a mechanism based on LRRK2 autophosphorylation. By mass photometry and Blue-Native PAGE, we could demonstrate that forced *in vitro* autophosphorylation leads to LRRK2 monomerization. We and others have previously shown that the Roc domain of LRRK2 is subject to autophosphorylation at multiple sites, which are in proximity to the dimerization interface within the Roc-COR module (Gloeckner et al., 2010; Greggio et al., 2009; Helton et al., 2021). Furthermore, we recently identified a novel intramolecular feedback regulation of the LRRK2 Roc G domain that depends on auto-phosphorylation of the G1+2 residue (T1343) in the Roc P-loop motif (Gilsbach et al., 2023). In contrast to wild-type, ATP dependent monomerization is abolished for T1343A, suggesting it plays a key role in regulating auto-phosphorylation dependent monomerization. Interestingly, in cells the T1343A mutant shows a similar increased Rab10-phosphorylation compared to the LRRK2 G2019S PD mutant, suggesting auto-phosphorylation-meditated regulation of the oligomerization state is an important step in the LRRK2 activation cycle.

In order to investigate the possible physiological role of an (auto)phosphorylation-induced monomerization, we focused on the interaction of LRRK2 with its substrates. In particular, changes in the oligomerization state have previously been associated with the activation of LRRK2 (Berger et al., 2010). For this purpose, we used our previously established integrative modelling approach relying on chemical crosslinking-derived constraints (Guaitoli et al., 2016). Interestingly, we could only see crosslinks between RAB10 and LRRK2 in the presence of GTP and RAB29. In addition, changes in the XL pattern between the LRRK2 apo and the Rab-bound state imply major conformational changes upon activation/switch to a substrate-binding competent state. These findings are in agreement with the recent tetrameric structure of RAB29-bound LRRK2 showing the kinase domain in an active conformation (Zhu et al., 2022). In contrast, a recently high-resolution structure of the LRRK2 dimer is considered to represents an inactive conformation of the kinase domain (Myasnikov et al., 2021). By integrative modelling, we could indeed show that less intermolecular crosslinks are satisfied when RAB10 is docked on the dimeric LRRK2. In contrast, the LRRK2 monomeric state gave models in better agreement with the experimental data, irrespective of the different simulation and prediction conditions employed. Interestingly, we also got more inter-molecular crosslinks satisfied when predicting the complex of monomeric state using AlphaFold-multimer, which was ran in an unbiased, crosslinking-agnostic way. In the latter case, we achieved more reliable complexes without using any available LRRK2 structural templates, again confirming that the FL APO conformation is not competent for RAB10 binding. The higher-scoring AF-multimer predicted interface of monomeric LRRK2 with RAB10 entails the α4 helix, which is also engaged in inter-molecular crosslinks with LRRK2 via Lys 136. Among the alternative docking poses predicted by AF-mulitmer, we also found conformations where RAB10’s T73 is highly proximal to the KIN catalytic pocket, further supporting the plausibility of these models (Supplementary Figure 3A). Notably, this interface is compatible a ternary complex between monomeric LRRK2-RAB10-RAB29 is predicted by AF-multimer (Supplementary Figure 3B). Intriguingly, the interface between Ras domain’s α4 helix, within the Rad domain C-terminal lobe, and a kinase domain has been already observed on homologues proteins (e.g. RHOE-ROCK1 complex, PDB: 2V55). Also, the predicted interfaces between RAB10 and LRRK2’s LRR and ARM domains are reminiscent of structures involving homologous Ras GTPases (e.g., RAN-RANBP1, PDB:1K5D and RHOA-SmgGDS, PDB:5ZHX, respectively). This suggests that AF-multimer is harnessing evolutionary and structural information of homologous proteins to predict the conformational changes required to dock RAB10 on LRRK2. In addition to the docking site on the ARM domain, another critical docking hotspot for RAB10 on LRRK2 is the LRR-Roc linker, which is predicted in contact with RAB10 as well as other Rab substrates (Supplementary Figure 3C), as previously suggested by MD simulations (Weng et al., 2023). This linker therefore appears to have a dual role: on one hand, it enables the conformational dynamics of LRRK2 necessary to unleash its kinase activity, as shown by MD simulation as well as by our hinge analysis on AF-multimer predicted conformers. On the other, it might contribute an additional docking site for Rab substrates binding. Our unsupervised, comparative analysis of monomeric LRRK2-RAB interfaces also suggests that activity modifier Rabs, such as RAB29, have a different ARM docking mode than Rab substrates (Supplementary Figure 3C), and that the differential ARM docking mode might dictate the functional role played by a given Rab interactor. Modeling analysis indicates that LRRK2 preferably binds and phosphorylates RAB10 in its monomeric state, suggesting distinct roles for LRRK2 oligomer species, where the dimeric form may be responsible for the GTPase activity whilst the LRRK2 monomer exerts kinase activities distinct from the oligomeric state. In fact, our data are implicating that the dimeric state favors auto-phosphorylation events while, in contrast, the monomeric form is the substrate-competent state acting as upstream-effector in a canonical sense. This is also in agreement with the observation that LRRK2 forms more monomers in presence of RAB29 in a Blue-Native gel analysis while no increase of higher-order oligomers is observed (Supplementary Fig. 1D). However, to understand the defined role of the LRRK2 monomeric state in substrate phosphorylation and the dynamics of kinase-active LRRK2 potentially shuttling between different oligomeric states at the membrane, certainly more work is needed.

In conclusion, we show the LRRK2-PD mutations located in different domains might be mediated by different pathophysiological mechanisms. More importantly, our study describes a novel feedback mechanism relaying on autophosphorylation induced monomerization by multifaceted approach. The integrative modelling indicates the LRRK2 monomer is the preferred conformation for binding and phosphorylating its physiological substrates, which might be the down-stream process of monomerization.

## Material and Methods

### Generation of LRRK2 expression constructs

The generation of N-terminal, SF-tagged (NSF), full-length LRRK2 expression constructs has been previously described (Gloeckner et al., 2010). N-terminal HA-FLAG RAB10 and RAB29 were generated as described in (Kishore et al., 2019).

### Cell culture and transfection

For expression of LRRK2, HEK293T cells (CRL-11268; American Type Culture Collection) were cultured in 14 cm dishes in DMEM (Sigma Aldrich) supplemented with 10 % (vol/vol) FBS (Sigma Aldrich) and 0.5 % (vol./vol.) Penicillin/Streptomycin antibiotics (Gibco). Cells were transfected at a confluence between 50 % and 70 % with 8 μg of plasmid DNA per dish using polyethyleneimine 25,000 MW (Polysciences) solution as described previously (Guaitoli et al., 2016). After transfection, cells were cultured for 48 h.

### Protein purification

Purification of (N)Strep-Flag-tagged LRRK2 as well as (N)HA-Flag RAB10 and RAB29 as well as RAB32 from 14 cm dishes (600 cm^2^) of confluent HEK293T cells was performed as previously described (Guaitoli et al., 2016). Briefly, after removal of the medium cells were lysed in 1 mL of lysis buffer (50 mM Hepes [pH 8.0], 150 mM NaCl, 1 mM DTT, 5 mM MgCl_2_, 5 % Glycerol, supplemented with 0.55 % Nonidet P40 substitute [Roche] and cOmplete^TM^ EDTA-free Protease Inhibitor Cocktail [Roche]), per culture dish. After cells were harvested and transferred to a Falcon tube, incubation in lysis buffer was carried on for 40 min at 4 °C under constant agitation. Cell debris and nuclei were removed by centrifugation at 10,000 × g and 4 °C for 10 min. Cleared lysates were incubated with 300 μL of settled ANTI-FLAG^TM^ M2 affinity resin (Sigma Aldrich) for 1 h at 4 °C. After incubation, the resin was transferred to microspin columns (GE Healthcare) and washed five times with 500 µl of washing buffer prior to final elution using 700 μL of elution buffer (Hepes-based buffer supplemented with 200 μg/mL Flag peptide). For mass photometry experiments, the elution buffer was supplemented by 100 μM G-nucleotide (either GppNHp or GDP).

### SDS and BN Gel Electrophoresis

SDS and Blue Native gel electrophoresis were performed as described in (Guaitoli et al., 2016). Briefly, SDS/PAGE was performed using 4–12% NuPage (Life Technologies) gradient gels. BN gel electrophoresis was performed on 4–12% NativePage gels (Life Technologies) according to the vendor’s protocols. The gels were stained with colloidal Coomassie.

### G-Nucleotide exchange

Nucleotide exchange of the G-proteins was performed a previously described (Lauer et al., 2019). Briefly, freshly purified LRRK2 protein was incubated in Elution buffer supplemented with 10 mM EDTA and a 10-fold molar excess of G-Nucleotide (either GDP or GTPγS) for at least 15 up to 30 minutes at 37 °C followed by an additional incubation on ice for 10 min. Finally, MgCl_2_ was added to the protein solution at a final concentration of 10 mM. The same protocol was used for nucleotide exchange for RAB10 and RAB29. For the CX-MS and the Blue Native gel analysis, the Rabs were loaded with either GTP or the non-hydrolysable analogue GTPγS.

### Thermal analysis measurements

Calorimetric measurements were carried out in HEPES-based purification buffer with a TA Instruments Nano-DSC (6300) equipped with capillary cells (sensitive volume of 0.3 mL) at 0.5°C·min^-1^ scan rate. A heating-cooling cycle followed by a second heating scan was scheduled in the temperature range from 5°C to 110°C for all experiments. A DSC with a capillary cell design was selected in order to prevent or at least mitigate protein aggregation phenomena (Pelosi et al., 2019).

Data were analyzed by means of the software THESEUS (Barone et al., 1992) following procedures reported in previous studies (Liggri et al., 2023; Mam et al., 2023). Briefly, the apparent molar heat capacity *C_P_(T)* of the sample was scaled with respect to the pre-denaturation region (namely, the region corresponding to *T* < *T_i_*, where *T_i_* is the highest temperature at which only the native protein is present, also indicated as molar heat capacity of the “native state”, *C_P,N_(T)*). The baseline subtraction provided the excess molar heat capacity *C_P_^exc^(T)* across the scanned temperature range. The area underlying the recorded peaks, so treated, directly corresponds to the relevant denaturation enthalpy, Δ*_d_H°.* The heat capacity drop, Δ*_d_C_P_*, across the signal was affected by a rather large error and therefore it was not considered in the present work. Furthermore, despite the use of a capillary nano-DSC, all the samples showed relevant aggregation phenomena, and the measurements were affected by high uncertainties as regards the observed enthalpy values. Nonetheless, a good reproducibility of the overall calorimetric profiles was obtained. Accordingly, here we only considered these intensive parameters (peak temperatures etc.), and the corresponding thermograms were reported by expressing the *C ^exc^* in terms of arbitrary units.

### Mass photometry

To evaluate the monomer/dimer ratio of LRRK2, mass photometry on a Refeyn Two^MP^ (Refeyn) instrument was performed (Chen et al., 2021). Prior to the experiment, the instrument was calibrated using a molecular weight standard (Native Marker, Invitrogen; diluted 1:200 in HEPES-based elution buffer). The LRRK2 protein was diluted to 2x of final concentrations (25, 50, 75, 100nM) used for the measurements. The optical focus was adjusted in 10μL elution buffer, before adding 10µl of diluted sample solution. After sufficient mixing of the samples, signals were recorded for 20s up to 1min depending on the obtained count numbers at different concentrations. The normalized relative dimer amount was determined using the following equation:

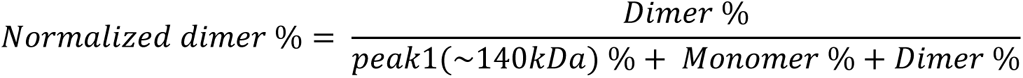

Three individual measurements were processed for each experimental condition, and data were analyzed by GraphPad Prism.

To induce autophosphorylation, LRRK2 was incubated with either 0.5 mM ATP or carrier (control) for 30 min at 30 °C. After dilution, the MP measurements were performed at a final protein concentration of 75 nM and 1min video recording. For MLi-2 treatment, LRRK2 was preincubated with MLi-2 or carrier (control) at a 10-fold molar access for 10 min on ice. Samples were then divided into two tubes and further incubated with 0.5 mM ATP or carrier (control) for 30 min at 30 °C.

### Chemical cross-linking mass spectrometry (CX-MS)

To study the LRRK2-Rab10 interaction, CX-MS was performed as previously described (Singh et al., 2022). Briefly, prior to the cross-linking reaction, a 1:5 complex of LRRK2 and RAB10+RAB29 was formed by incubation in elution buffer for 30 minutes at 4 °C under constant mixing conditions. For subsequent NHS-ester–based chemical cross-linking, the CID-cleavable crosslinker DSSO (Thermo Fisher, 12.5 mM stock solution in DMSO) was added to approximately 150 μg of the preformed LRRK2-RAB10 complex, in a total volume of approximately 0.4 mL of elution buffer to reach a final ratio in terms of DSSO to protein of 50:1. The reaction was carried out under constant mixing conditions for up to 45 min, then stopped by adding Tris⋅HCl (pH 7.5) solution to a final concentration of 100 mM. The mix was incubated for 15 min at room temperature under constant shaking, following precipitation by chloroform-methanol. Proteolysis was performed dissolving the protein precipitates in 50 mM ammonium bicarbonate containing 0.2% RapiGest (Waters) reduced with DTT and alkylated with iodoacetamide, and proteolysis was performed by adding either 2 μg of trypsin (Promega) or a combination of 2 μg of trypsin and 0.2 μg of LysC (Promega). Samples were incubated at 37 °C overnight. After proteolysis, the RapiGest surfactant was hydrolyzed by acidification and removed by centrifugation. The samples were further cleaned up via C18-StageTips (Thermo Fisher) following standard protocols. Sample volumes were reduced to ∼10 μL in a SpeedVac. Subsequently, SpeedVac-dried samples were re-dissolved in 30 μL of SEC buffer [30% (vol/vol) acetonitrile, 0.1% TFA]. Samples were loaded in two fractions (15 μL) and separated on a Superdex Peptide PC 3.2/30 column (GE-Healthcare) at a flow rate of 50 μL on an Äkta pure LC system (Cytiva). One hundred-microliter fractions were collected. The fractions were subsequently vacuum-dried and re-dissolved in 5% TFA prior to LC-MS analysis on an Orbitrap Fusion instrument (Thermo Scientific) using an MS2/MS3 scheme implemented in the instrument control software. The resulting RAW data were analyzed by the Proteome Discoverer Software (v2.4.1.15).

### Integrative modeling

We predicted the 3D structure of full length LRRK2, either monomeric or dimeric, in complex with RAB10, by using cross-linking/MS-derived distance restrained docking calculations through the Integrative Modeling Platform (IMP) (Webb et al., 2018) package, release 2.17.0. We employed the Python Modeling Interface (PMI), adapting the scripted pipeline of a previously described procedure (Chen et al., 2021) and consisting in the following key steps: (1) gathering of data, (2) representation of domains and/or subunits and translation of the data into spatial restraints, (3) configurational sampling to produce an ensemble of models that optimally satisfies the restraints, and (4) analysis and assessment of the ensemble.

### System representation

The monomeric and dimeric LRRK2 structures (PDB ID: 7LHW and 7LHT, respectively), as well as the RAB10 structure (PDB ID: 5SZJ), were used as input structures for IMP calculations. LRRK2 and RAB10 were represented by beads arranged into either a rigid body or a flexible string. The beads representing a structured region were kept rigid with respect to one another during configurational sampling (i.e. rigid bodies). Based on the inspection of experimental as well as predicted (i.e. https://alphafold.ebi.ac.uk/entry/Q5S007) structures, we defined as flexible beads the following regions: aa 1328-1329, corresponding to the LRR-Roc linker, and aa 1657-1667, corresponding to the CORA-CORB linker.

Every other region was defined as a rigid-body. For both the monomeric and dimeric LRRK2-RAB10 complexes, we set up two simulations: one where only the LRR-Roc linker was kept flexible, and another where both LRR-Roc and CORA-CORB were kept flexible.

### Bayesian scoring function

The cross-linking data were encoded into a Bayesian scoring function that restrained the distances spanned by the cross-linked residues (Erzberger et al., 2014). The Bayesian approach estimates the probability of a model, given information available about the system, including both prior knowledge and newly acquired experimental data (Erzberger et al., 2014). Briefly, using Bayes’ theorem, we estimate the posterior probability p(M D,I), given data D and prior knowledge I, as p(M D, I) ∝ p(D M, I)p(M, I), where the likelihood function p(D M,I) is the probability of observing data D, given I and M, and the prior is the probability of model M, given I. To define the likelihood function, one needs a forward model that predicts the data point (i.e. the presence of a cross-link between two given residues) given any model M and a noise model that specifies the distribution of the deviation between the observed and predicted data points. To account for the presence of noisy cross-links, we parameterized the likelihood with a set of variables {ψ} defined as the uncertainties of observing the cross-links in a given model (Erzberger et al., 2014; Robinson et al., 2015). A distance threshold of 20 Å was employed to model DSSO cross-linkers.

### Sampling model configurations

Structural models were obtained by Replica Exchange Gibbs sampling, based on Metropolis Monte Carlo sampling (Rieping et al., 2005). This sampling was used to generate configurations of the system as well as values for the uncertainty parameters. The Monte Carlo moves included random translation and rotation of rigid bodies (4 Å and 0.03 rad, maximum, respectively), random translation of individual beads in the flexible segments (5 Å maximum), and a Gaussian perturbation of the uncertainty parameters. A total of 1M models per system were generated, starting from 100 random initial configurations and sampling with temperatures ranging between 1.0 and 2.5.

### Analysis of the model ensemble

The 200 best scoring models (i.e. solutions) for each docking run were clustered to yield the most representative conformations. For each ensemble, the solutions were grouped by k-means clustering on the basis of the r.m.s. deviation of the domains after the superposition of LRRK2. The precision of a cluster was calculated as the average r.m.s. deviation with respect to the cluster center (i.e. the solution with the lowest r.m.s. deviation with respect to the others).

For each cluster, we calculated the number of satisfied MS/cross-links by measuring the number of Cα pairs, corresponding to cross-linked Lys, whose distance was shorter than 35 Å. We recorded the fraction of satisfied cross-links, given by the number of satisfied cross-links over the total, for all the cluster members. We finally reported the fraction of satisfied cross-links for the best scoring solution, as well as the maximum fraction obtained for a single cluster conformer and the aggregate fraction obtained by considering all the cluster members together.

For visualization purposes, we generated all atoms models by starting from the Cα traces generated from IMP and using the automodel() function from Modeler (Sali and Blundell, 1993).

### AlphaFold-multimer predictions

We used Alphafold-multimer v2.3.1, 3D models of the human LRRK2-RAB10 complex. The necessary databases for running AlphaFold-multimer were downloaded on 12 January 2023 (Liu et al., 2023).

To encourage the exploration of a broader conformational space and avoid being biased by experimental templates, we generated the models without the utilization of structural templates. This was done by setting the flag –max template date=01-01-1900 to ensure that AlphaFold-multimer did not rely on any experimental structural data.

In the following analysis we focused on the best model out of the 25 generated. The evaluation of the models was based on the default scoring system used by AlphaFold-multimer (0.2*pTM + 0.8*ipTM). We generated AF-multimer complexes of monomeric and dimeric LRRK2 in complex with RAB10, as well as of monomeric LRRK2 in complex with both RAB10 and RAB29.

We also predicted the 3D complexes of monomeric LRRK2 with Rab GTPases reported to interact with LRRK2 in IntAct (Porras et al., 2020). We performed analysis of 3D interfaces using our previously established protocol (Matic et al., 2023), through which we considered residue-residue contacts as those having the Cβ spatially closer than 8 Å (Cɑ for glycine). We generated a contact fingerprint where each Rab interactor is described by a vector, whose index corresponds to LRRK2 positions, and numerical values to the number of predicted complexes involving a given Rab showing that contact. We employed the contact fingerprint to unsupervised cluster LRRK2-RAB complexes on the basis of their interface structural characteristics.

### Structural analysis

The mapping of chemical crosslinks to structural models was performed through a custom script. We computed the distance between the Cɑ coordinates of the crosslinked residues and we considered the crosslink satisfied if the distance was smaller than 35 Å.

To identify the structural rearrangements of the models with respect to the monomeric experimental structure (PDB:7li4) we used DynDom6D (Veevers and Hayward, 2019) (v1.0, setting the grid size to 8 Å, the block factor to 3 and the minimum domain size to 100).

We employed Pymol (v2.4.1) and ChimeraX (v1.5) to generate 3D cartoon representations. We employed customized scripts in python (version 3.8.11), using matplotlib (v3.6.0), seaborn (v0.11.1), and biopython (v1.78) libraries to generate statistical analysis and plots. We calculated residue-residue contact by using a customized script derived from the CIFPARSE-OBJ C + + library (https://mmcif.wwpdb.org/docs/sw-examples/cpp/html/index.html).

## Supporting information

Supplemental information on the DSC analysis

## Acknowledgements

The authors wish to thank the staff of the Core Facility for Medical Bioanalytics (Tübingen), especially Franziska Klose, for technical assistance. The work was supported by The Michael J. Fox Foundation for Parkinson’s Research (grant No. 8068.04).

## Author Contributions

G.G., F.S. and M.S. performed nDSC experiments. G.G, X.Z., L.M.S. and K.T. performed the mass photometry measurements. F.v.Z. and G.G. generated expression constructs and performed protein purifications. G.G., F.S., D.F., X.Z., F.Y.H., A.K. and C.J.G. analyzed data. P.M. and F.R. performed the computational modelling. G.G., F.R, A.K. and C.J.G. designed the study. G.G, F.S., X.Z., D.F., F.R., A.K. and C.J.G. wrote the manuscript.

## Data availability

Mass photometry raw data to generate Figure 3 have been deposited on Zenodo (DOI: 10.5281/zenodo.8387721). Mass spectrometric (CX-MS) RAW data have been deposited on PRIDE (Acc. No. PXD044138).

## Abbreviations

AF: AlphaFold
Ct: *Chlorobium tepidum,*
iPD: idiopathic PD
MP: mass photometry
nDSC: nano differential scanning calorimetry
PD: Parkinson’s disease
RCKW: Roc-COR-kinase-WD40 LRRK2 construct
SF: tandem Strep/ FLAG tag
CX-MS: chemical crosslinking (CX) combined with Mass-spectrometry-based (MS) mapping.

## Supplemental Figures and Tables

**Supplementary Figure 1:**
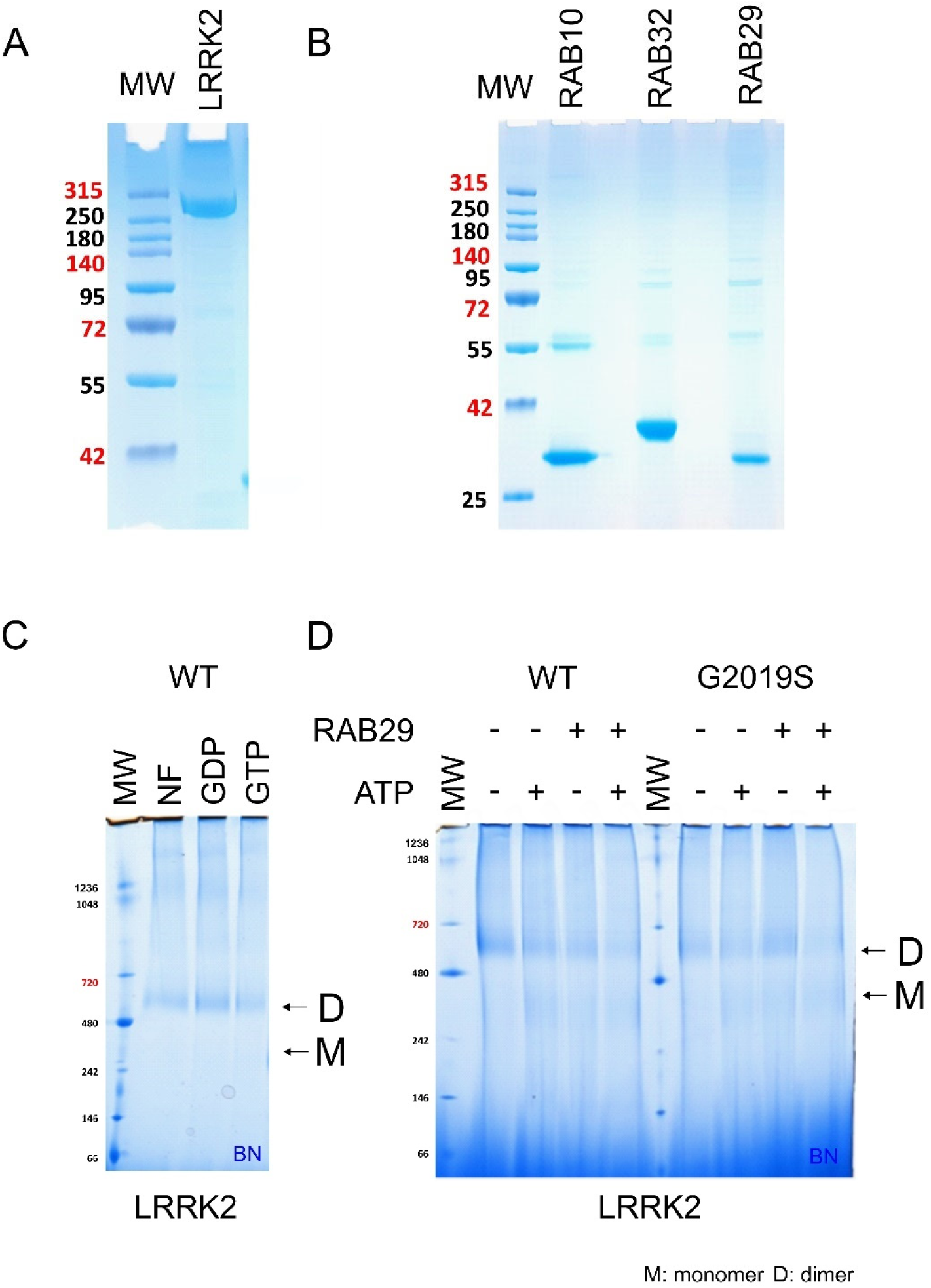
(A-B) Purification of LRRK2 and Rab proteins. Representative SDS PAGE gels stained with colloidal Coomassie are shown for (A) SF-tagged LRRK2 and (B) FLAG-HA-tagged Rab proteins. LRRK2 was either purified via the tandem Strep-tag II or via the FLAG tag. Rab proteins were purified via the FLAG-tag. (C-D) Blue Native analysis of LRRK2. (C) Nucleotide dependence of LRRK2 oligomerization (NF: nucleotide-free, GDP, GTP). (D) ATP-incubation (5mM) induced monomerization of LRRK2 WT and G2019S. In addition, the influence of RAB29 on the LRRK2 oligomeric state was tested. A smaller part of the same gel is shown in Figure2B (lanes 1 and 2/ LRRK2 WT +/- ATP).

**Supplementary Figure 2:**
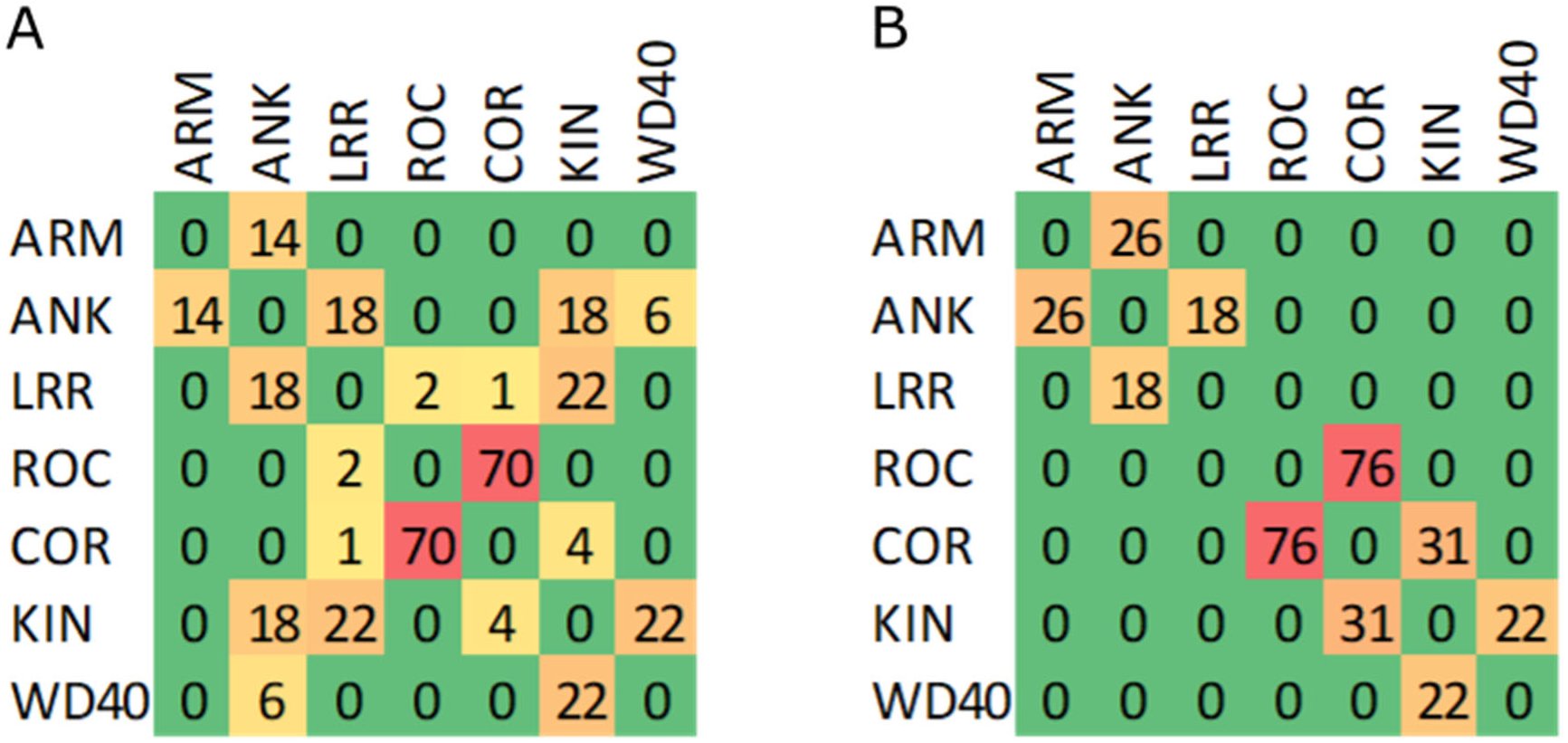
LRRK2 domain contact analysis of (A) LRRK2 FL APO structure (PDB:7LHW) and (B) full-length LRRK2 predicted by AlphaFold2 (AF-Q5S007-F1).

**Supplementary Figure 3:**
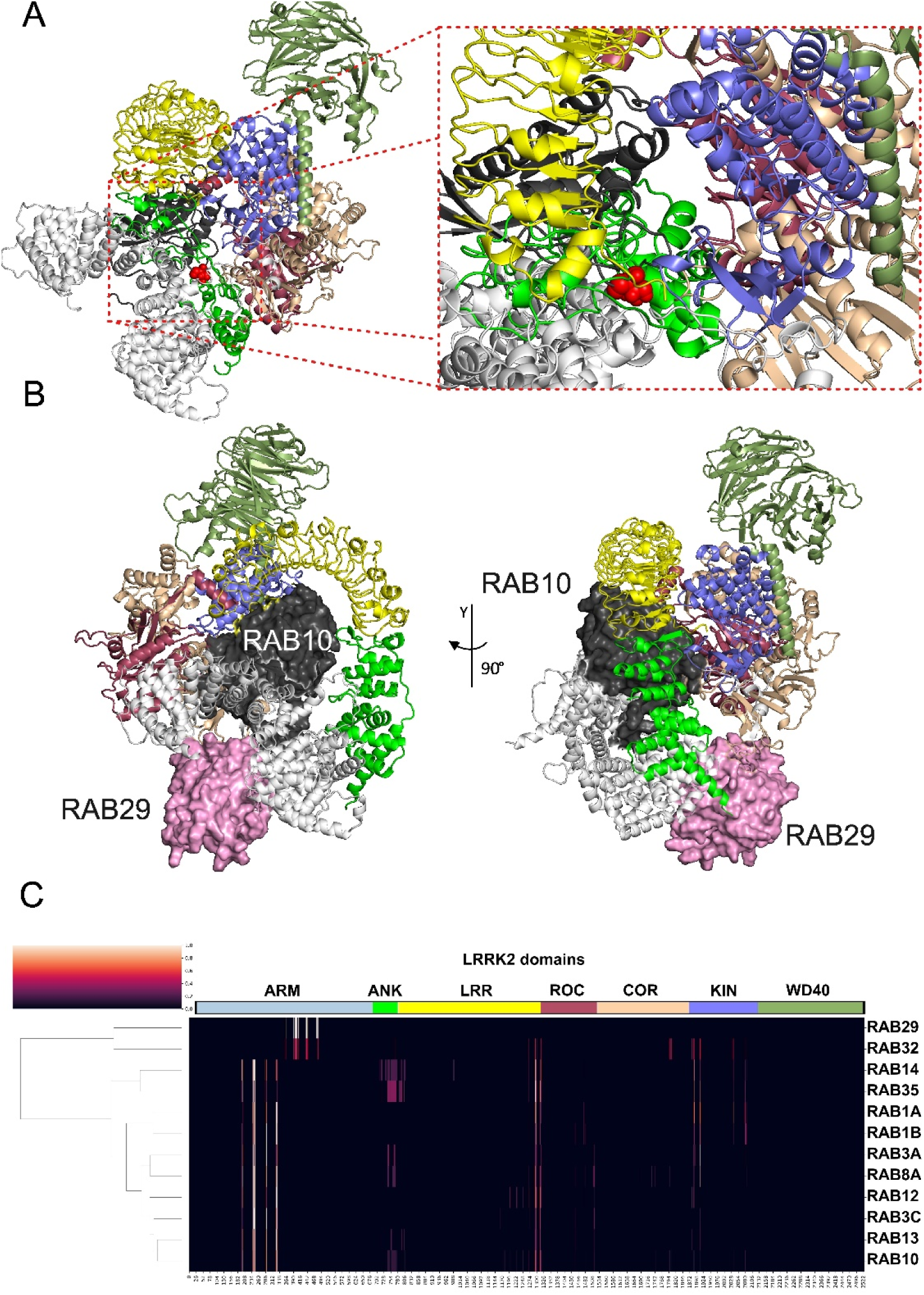
(A) alternative docking mode of RAB10 on monomeric LRRK2 predicted by AF-multimer (without templates) showing T73 phosphosite (indicated as red spheres) close to the Kinase domain catalytic pocket. Cartoon coloring scheme is domain specific. (B) Model of monomeric LRRK2 bound to RAB29 and RAB10 predicted by AF-Multimer. (C) Interface contact sites of AF-multimer predicted complexes between monomeric LRRK2 and RAB proteins previously shown to interact with LRRK2 according to the IntAct database. The heatmap shows the normalized number of contacts seen across the predicted models for each interacting RAB (y axis) for each LRRK2 position (x axis).

**Supplementary Figure 4:**
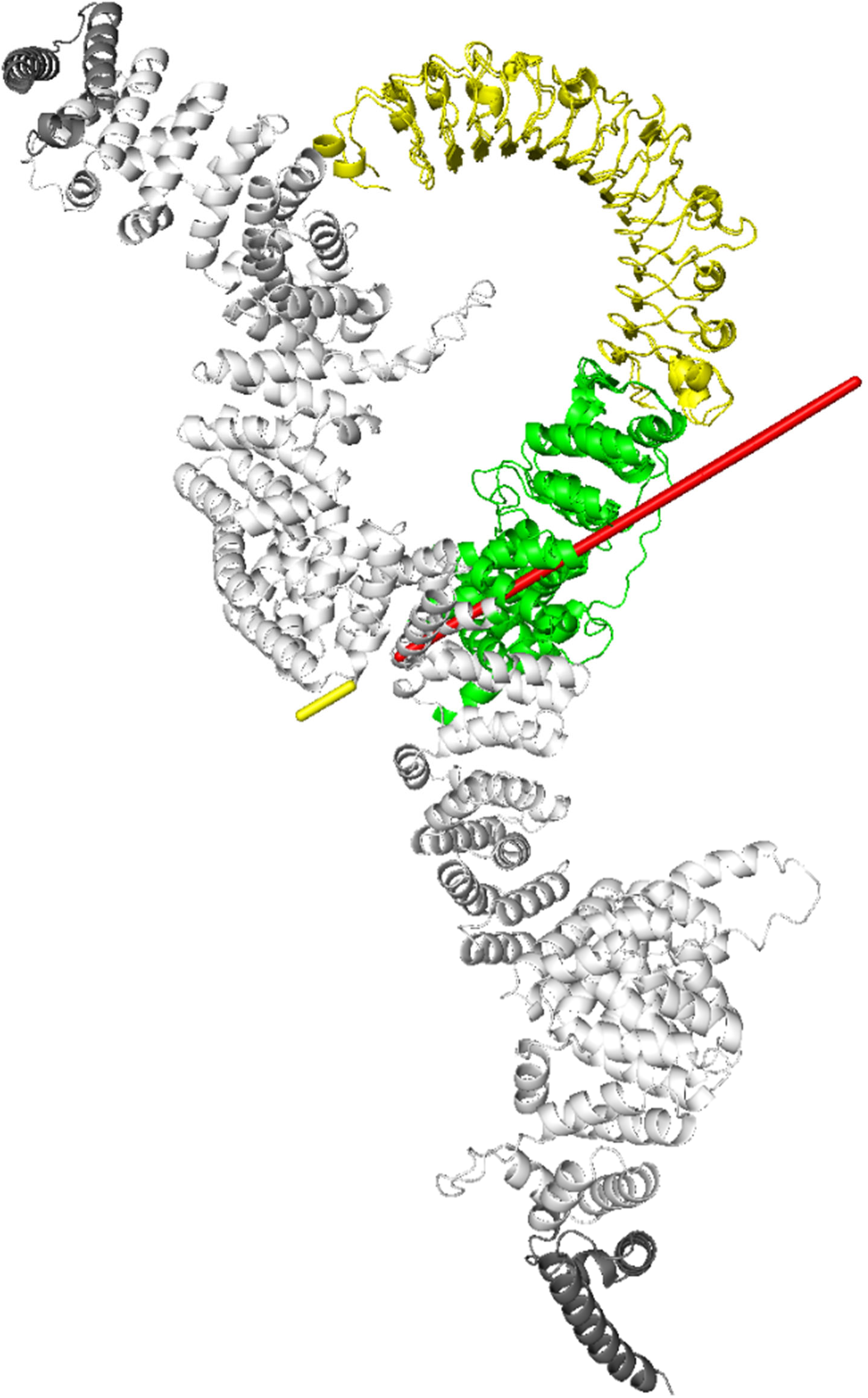
Superimposition of the N-terminal domains, i.e. LRR (yellow), ANK (green) and ARM (white), of LRRK2 structure in the full-length APO state (predicted by AF2 using available experimental structural templates) with the structure predicted by AF-multimer in complex with RAB10. Red axis indicates the conformational change identified between the ANK and ARM domains.

## Notes

https://doi.org/10.5281/zenodo.8387721

### Competing Interest Statement

The authors have declared no competing interest.

### Summary of Updates

- textual changes throughout the manuscript - additional supplementary data were added

