## Supplemental information on the DSC analysis for "Biophysical analysis reveals autophosphorylation as an important negative regulator of LRRK2 dimerization"

### *Thermodynamic analysis on the LRRK2 apo-protein n-DSC data*

We mentioned in the main text that the n-DSC measurements for the LRRK2 apo-protein (Fig. 1 in the main text) were affected by aggregation phenomena observed at high temperatures, introducing a considerable experimental error on the evaluation of the overall experimental denaturation enthalpy $\Delta_d H^{exp}$, and preventing us from performing a straight thermodynamic analysis in absolute enthalpic and entropic terms. Nonetheless, the calorimetric profiles are still highly informative and allowed at least a qualitative assessment of the overall denaturation mechanism.

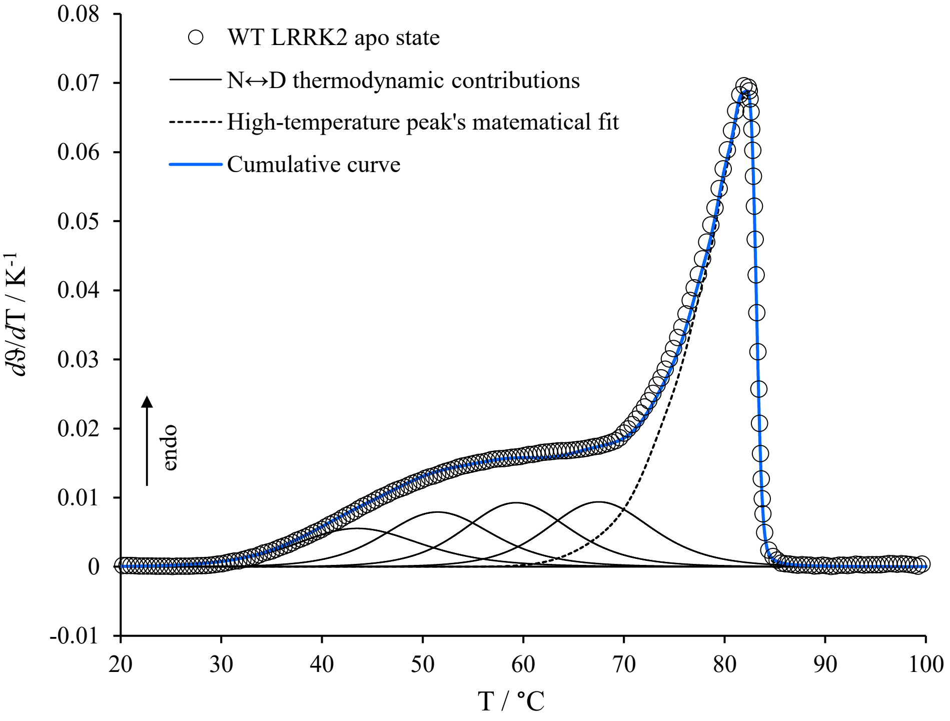


***Fig. S1.*** *Normalized nano-DSC thermogram of the purified full-length LRRK2 apo-state (circles) and an example of tentative fit of only the first part of the thermogram (below 70°C) through the application of a multi-domain single-step native-to-denatured state equilibrium mechanism (black solid curves). The remaining part of the thermogram (above 70°C) was obtained by a mere mathematical fitting in order to tentatively isolate the late calorimetric events (black dashed curve). The sum of both the thermodynamic contributions and the high-temperature mathematical curve is also reported as a cumulative curve in blue.*

To this purpose, we considered the extent of reaction $\vartheta(T)$ for our experimental calorimetric profile

$$\vartheta(T) = \frac{\int_{T_0}^{T} C_p^{exc}\,\mathrm{d}\,T}{\Delta_d H^{exp}} \tag{1}$$

i.e., a function ranging from 0 to 1 that reflects the fraction of molecules that have undergone the denaturation at a given temperature. The temperature derivative of such a function, $d\vartheta/dT$, reported in Fig. S1, just corresponds to the normalized DSC thermogram and was considered for the tentative application of various thermodynamic models. Indeed, the function *ϑ(T)* only depends on the calorimetric profile, whereas it does not depend on the overall experimental enthalpy, which instead becomes a mere normalization factor.

At first, we considered the application of the simplest thermodynamic model, which complains the contribution of more protein domains *i*, each undergoing denaturation independently according to a single-step native-to-denatured state equilibrium mechanism (Fessas et al., 2001)

$$N_i \overset{K}{\leftrightarrow} D_i \qquad (2)$$

by using the thermodynamic equations deriving from model (2), i.e.,

$$\frac{\mathrm{d}\vartheta_i}{\mathrm{d}T} = \frac{\Delta_d H_i^{\circ 2}}{RT^2} \frac{K_i(T)}{[1 + K_i(T)]^2} \frac{1}{\Delta_d H^{exp}} \qquad (3)$$

where

$$K_i(T) = \exp\left[-\frac{\Delta_d H_i^{\circ}}{R}\left(\frac{1}{T} - \frac{1}{T_{d,i}}\right)\right] \qquad (4)$$

whereas $\Delta_d H_i^{\circ}$and $T_{d,i}$ are the denaturation enthalpy and the denaturation temperature, respectively, for each independent domain.

The results of these attempts indicated that the first part of the thermogram can be described as a sum of more protein domains each undergoing denaturation independently according to (2), whereas such an approach was clearly unable to describe the high-temperature region of the curve.

Accordingly, by using a simple mathematical curve to take into consideration the high-temperature region, an example of such a tentative to fit the first part of the thermogram is reported in Fig. 1S. We would like to highlight here that, though the thermodynamic curves shown in Fig. S1 are normalized (i.e., divided by the overall experimental enthalpy), the denaturation enthalpy $\Delta_d H_i^{\circ}$ of each deconvoluted domain is included into the respective curve equation, as shown in equations (3) and (4), and the curve profile peculiarities are hence strongly dependent on $\Delta_d H_i^{\circ}$ values (the higher the $\Delta_d H_i^{\circ}$ the higher the height/width ratio of the thermodynamic curve). Nonetheless, when in the presence of uncertainties on the overall experimental profile, a straight validation of the $\Delta_d H_i^{\circ}$ parameter for each domain obtained from this kind of best fit cannot be performed. Furthermore, as regards the number of the thermodynamic contribution, degeneracy cannot be excluded (i.e., depending on the mathematical curve chosen for the description of the high-temperature region, we may not discriminate between different sets of theoretical solutions that may also fit the first part of the thermogram). For this reason, we avoid to report the fitting parameters ($\Delta_d H_i^\circ$ and $T_{d,i}$) and we only limit to state that the first part of the thermogram is compatible with such a type of scenario.

Conversely, according to the literature (Ausili et al., 2013; Fessas et al., 2001), the thermogram portion above 70°C reveals a calorimetric profile that is typical of a denaturation mechanism that complains a dissociation process of a multimer (left tail asymmetry) concomitant with its denaturation (Privalov and Potekhin, 1986), i.e.,

$$N_n \overset{K}{\leftrightarrow} nD \ . \tag{5}$$

being *n* the dissociation stoichiometry

The best fitting attempt applied to the second part of the thermogram (dashed curve in Fig. S1) according to this model resulted to be close to the experimental profile by using *n*=2, but still unsatisfactory probably because of the irreversible aggregation process that is clearly enhanced within this high-temperature region. Accordingly, basing on the Lumry-Eyring models (Sanchez-Ruiz, 1992), we included a post-denaturation step in our thermodynamic model, i.e.,

$$N_n \overset{K}{\leftrightarrow} nD \xrightarrow{k_{app}} nA \tag{6}$$

where the form *A* indicates the aggregated state.

According to model (6), the theoretical trace is obtained through the equation (Sanchez-Ruiz, 1992)

$$\frac{\mathrm{d}\vartheta}{\mathrm{d}T} = \frac{E_{app}}{RT_{max}^2} \exp\left(\frac{E_{app}\Delta T}{RT_{max}^2}\right)\left[1 + \frac{1-n}{n}\exp\left(\frac{E_{app}\Delta T}{RT_{max}^2}\right)\right]^{\frac{1}{n-1}} \tag{7}$$

where:

- *n* is the dissociation stoichiometry;
- $E_{app} = E_a + \Delta_d H^\circ$, with $\Delta_d H^\circ$ as the overall equilibrium transition enthalpy expressed per monomer (i.e., the denaturation enthalpy including a potential enthalpic contribution given by the dissociation) and $E_a$ as an activation energy to account the kinetic step of aggregation $D \rightarrow A$ by using a first order kinetic constant $k_{app}$, whose temperature dependence follows the Arrhenius equation;

- $\Delta T = (T - T^{*})$, that is the difference between the variable $T$ and a reference temperature at which the aggregation rate corresponds to $k_{app}$=1 min$^{-1}$.

The best fitting attempt achieved according to this model, shown in Fig. S2, resulted to be rather satisfying for the second part of the DSC curve and was obtained again by assuming $n$=2.

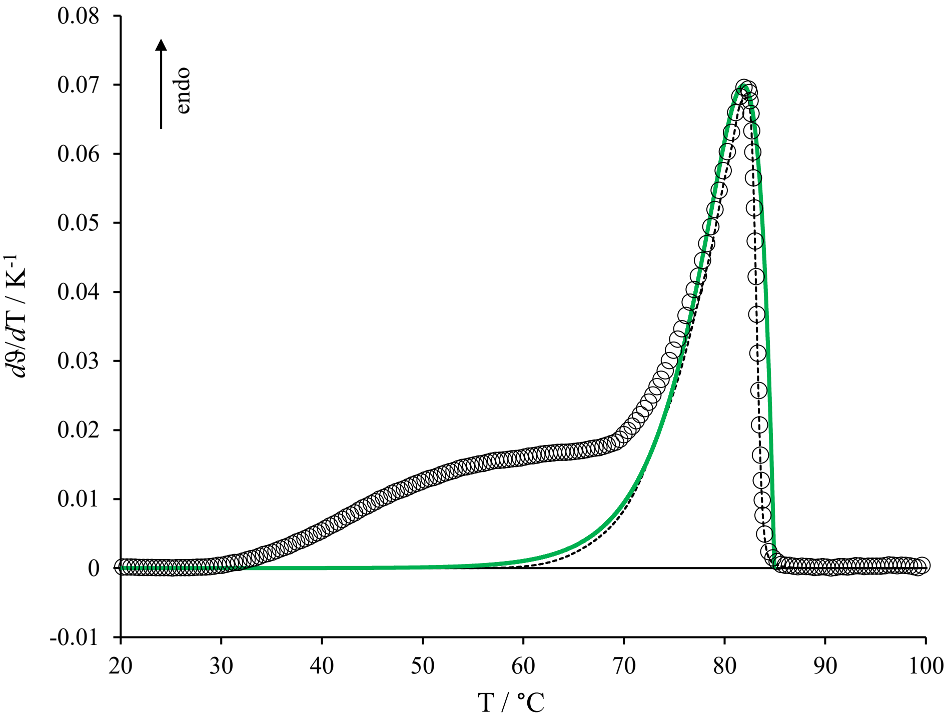


***Fig. S2.*** *Thermodynamic deconvolution of the late part of the LRRK2 apo-state thermogram through the application a model that complains a dimer dissociation process concomitant with its denaturation and followed by aggregation phenomena (green curve). The LRRK2 overall experimental profile (circles) and the mathematical curve representing the late calorimetric events (black dashed trace) are also reported for comparison.*

In conclusion, being aware of all the uncertainties as regards the absolute enthalpic values, the thermodynamic analysis of the n-DSC data indicated an overall denaturation mechanism of the LRRK2 apo-protein that complains the single-step independent denaturation of the low-stability domains followed by a dissociation process concomitant with the denaturation of the most stable domains as shown, for the sake of completeness, by the overall fitting attempt in Fig. S3.

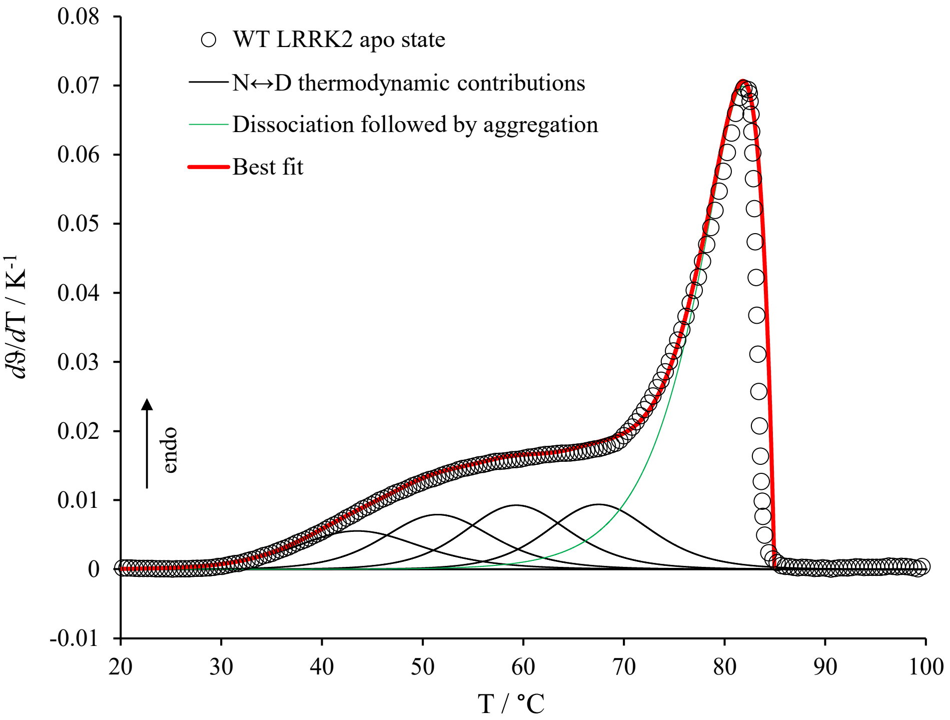


***Fig. S3.*** *Overall representation of the nano-DSC thermogram of LRRK2 apo-state (circles) together with a tentative complete thermodynamic deconvolution of the profile. The domains compatible with a single-step native-to-denatured state equilibrium mechanism are shown as black curves; the contribution compatible with a dimer dissociation process concomitant with its denaturation and followed by aggregation phenomena is shown as a green curve; the overall best-fit curve is shown as a red curve.*

A similar conclusion concerning the overall denaturation mechanism can be also drawn for the system LRRK2+GDP, which, among the peculiarities reported in the main text, exhibits a very similar denaturation profile.

Technical note: The fit attempts based on the denaturation thermodynamic models were accomplished using the nonlinear Levenberg – Marquardt method (Press et al., 2007).

***Supplementary References***


Ausili, A., Pennacchio, A., Staiano, M., Dattelbaum, J.D., Fessas, D., Schiraldi, A., and D'Auria, S. (2013). Amino acid transport in thermophiles: characterization of an arginine-binding protein from Thermotoga maritima. 3. Conformational dynamics and stability. *J Photochem Photobiol B 118*, 66-73 doi: 10.1016/j.jphotobiol.2012.11.004.

Fessas, D., Iametti, S., Schiraldi, A., and Bonomi, F. (2001). Thermal unfolding of monomeric and dimeric beta-lactoglobulins. *Eur J Biochem 268*, 5439-5448 doi: 10.1046/j.0014-2956.2001.02484.x.

Press, W.H., Teukolsky, S.A., Vetterling, W.T., and Flannery, B.P. (2007). Nonlinear Models. In NUMERICAL RECIPES Third Edition-Numerical Recipes: The Art of Scientific Computing, Third Edition, p. 799.

Privalov, P.L., and Potekhin, S.A. (1986). Scanning microcalorimetry in studying temperature-induced changes in proteins. *Methods in enzymology 131*, 4-51 doi: 10.1016/0076-6879(86)31033-4.

Sanchez-Ruiz, J.M. (1992). Theoretical analysis of Lumry-Eyring models in differential scanning calorimetry. *Biophys J 61*, 921-935 doi: 10.1016/S0006-3495(92)81899-4.
